# The subcortical correlates of self-reported sleep quality

**DOI:** 10.1101/2024.05.29.596530

**Authors:** Martin M. Monti

## Abstract

**Study objectives:** To assess the association between self-reported measures of sleep quality and cortical and subcortical local morphometry.

**Methods:** Sleep quality, operationalized with the Pittsburgh Sleep Quality Index (PSQI), and neuroanatomical data from the full release of the young adult Human Connectome Project dataset were analyzed (N=1,112; 46% female; mean age: 28.8 years old). Local cortical and subcortical morphometry was measured with subject-specific segmentations resulting in voxelwise gray matter difference (i.e., voxel based morephometry) measurements for cortex and local shape measurements for subcortical regions. Associations between the total score of PSQI, two statistical groupings of its subcomponents (obtained with a principal component analysis), and their interaction with demographic (i.e., sex, age, handedness, years of education) and biometric (i.e., BMI) variables were assessed using a general linear model and a nonparametric permutation approach.

**Results:** Sleep quality-related variance was significantly associated with subcortical morphometry, particularly in the bilateral caudate, putamen, and left pallidum, where smaller shape measures correlated with worse sleep quality. Notably, these associations were independent of demographic and biometric factors. In contrast, cortical morphometry, along with additional subcortical sites, showed no direct associations with sleep quality but demonstrated interactions with demographic and biometric variables.

**Conclusions:** This study reveals a specific link between self-reported sleep quality and subcortical morphometry, particularly within the striatum and pallidum, reinforcing the role of these regions in sleep regulation. These findings underscore the importance of considering subcortical morphology in sleep research and highlight potential neuromodulatory targets for sleep-related interventions.

**Statement of Significance:** This study provides evidence for the role of subcortical structures in sleep regulation, demonstrating that poorer self-reported sleep quality is associated with reduced shape measures in the bilateral caudate, putamen, and left pallidum. These associations were independent of demographic and biometric factors, distinguishing subcortical morphometry as a unique correlate of sleep quality. In contrast, cortical morphology exhibited no direct relationship with sleep quality but showed interactions with factors such as sex, BMI, and age, highlighting the complexity of brain-sleep relationships. These findings contribute to a growing body of literature emphasizing the basal ganglia’s role in non-motor functions, including sleep regulation, and suggest potential neuromodulatory targets for sleep-related interventions, such as transcranial focused ultrasound or deep brain stimulation.

## Introduction

Highly conserved across the animal kingdom [1], sleep is important for cognitive, social, and emotional functioning [2–4] across the lifespan [5, 6]. Indeed, sleep deprivation has been associated with cognitive and emotional dysfunction [7–12], sleep disturbances are associated with psychiatric illnesses [13– 16] and are part of the known non-motor symptomatology of multiple neurodegenerative disorders [17–22].

Despite the importance of sleep, our understanding of its relationship with brain morphology remains in the early phases [23, 24], and has often returned conflicting findings. Poor sleep quality, in the context of insomnia disorder, for example, has been associated with morphometric differences (e.g., thinner cortex and/or smaller volumes) in orbitofrontal, mesial, and superior frontal cortices, precuneus, hippocampus, thalamus [25–29], although with substantial variance and inconsistencies across studies [30–35], potentially due to methodological differences and relatively small samples. Self-reported sleep quality in healthy individuals, across the lifespan, has been associated cross-sectionally with a number of regions, including frontal and parietal cortices, hippocampus, amygdala, and thalamus [36–38], and longitudinally with changes spanning multiple cortices (i.e., frontal, temporal, and parietal) [36, 39] including hippocampus [40] and the posterior cingulate cortex [41] specifically (see 42] for a recent overview of findings). Further conflating the issue, recent work uncovered moderating effects of several variables including sex [34], age (at least during adolescence) [43], chronotype [44], and developmental socioeconomic context [45] on the neural correlates of sleep quality. Even as large sample studies emerge [34, 39, 40, 46– 50], results are yet to coalesce and remain variable, perhaps also due also to different methodological choices [51], and are often associated with relatively small effect sizes. Nonetheless, understanding the relationship between brain and sleep is key to the development of novel interventions aimed at mitigating sleep-related disorders [52].

In what follows, the full release of the Human Connectome Project (HCP) [53] is leveraged to assess the relationship between self-reported sleep quality measures [54] and brain morphometry using local (i.e., voxelwise) cortical [55, 56] and subcortical [57] measures, as opposed to the global and regional features (i.e., average regional volumes, areas, and thickness) used in some prior work [40, 48, 50]. In particular, while prior studies have leveraged a subset of the same data for assessing functional and structural associations with sleep quality [34, 58], the approach used here adopts a more sensitive (i.e., voxelwise) approach at for both cortex, assessing differences in local concentration of gray matter [55], and subcortex, assessing differences in local shape of deep brain nuclei [57]. Importantly, these voxelwise methods have previously been shown to be more sensitive and better able to capture differences between groups than more global (e.g., regional) measures [59, 60].

As described below, this sensitive approach uncovers systematic associations between self-reported sleep quality and subcortical shape. In addition, variance in cortical and subcortical neuroanatomy is also associated with the interaction of sleep quality related measures and demographic and biometric variables, highlighting the multifaceted nature of brain-sleep relationships.

## Methods

### Sample description

This project is based on the Q3 release of the Human Connectome Project (HCP [53]). Out of the 1206 total participants, 93 could not be included in the analysis due to failure to acquire at least one T1-weighted MRI, resulting in an included sample of 1,113 observations. (As described below, one additional observation was excluded due segmentation failure, resulting in a final analyzed sample of 1,112 participants). The final analyzed sample included data from 605 female and 507 male participants, with a median age of 29 years. As described in the statistical analysis section, further demographic (i.e., handedness, years of education) and biometric (i.e., body mass index; BMI) factors, were also collected and used as covariates in assessing the relationshp between self-reported sleep quality and neuroanatomy (See Tab. 1 and Tab. SOM1). Because BMI was not recorded for one participant, and years of education was not recorded for two (different) participants, data was imputed with a median imputation strategy, which has been previously been shown to be equally effective as more complex approaches [61], particularly in datasets with low degrees of data missingness [62]. While this approach could decrease covariate variability, the standard deviation for each variable is only decreased by 0.05% (i.e., from 1.799 to 1.798) and 0.04% (i.e., from 5.199 to 5.197) for years of education and BMI, respectively. This is consistent with the fact that less than 0.2% of the data for each variable is being imputed (in fact, 0.18% and 0.09% for years of education and BMI, respectively).

**Table 1.** Sample demographic, biometric, and PSQI information. (Abbreviations: Med., median; sd., standard deviation; PC1, PSQI Principal Component 1; PC2, PSQI Principal Component 2; Subj. sleep qual., subjective sleep quality; sleep dur., sleep duration; sleep eff., sleep efficiency; sleep ons. lat., sleep onset latency; sleep disturb., sleep disturbances; sleep med., use of sleep medications; daytime dysf., daytime dysfunction;. See Tab. SOM1 for the same information broken down by sex.)

| Variables | Med | Mean | sd | Range |
| --- | --- | --- | --- | --- |
| PSQI Total Score | 4 | 4.804 | 2.759 | [0, 19] |
| Subj. sleep qual. (PC1) | 1 | 0.894 | 0.635 | [0,3] |
| Sleep dur. (PC1) | 0 | 0.569 | 0.815 | [0,3] |
| Sleep eff. (PC1) | 0 | 0.456 | 0.795 | [0,3] |
| Sleep ons. lat. (PC2) | 1 | 0.967 | 0.817 | [0,3] |
| Sleep disturb. (PC2) | 1 | 1.093 | 0.485 | [0,3] |
| Sleep med. (PC2) | 0 | 0.237 | 0.677 | [0,3] |
| Daytime dysf. (-) | 1 | 0.588 | 0.635 | [0,3] |
| Age | 29 | 28.799 | 3.696 | [22,37] |
| Handedness | 80 | 65.845 | 44.208 | [-100,100] |
| Education | 16 | 14.927 | 1.799 | [11,17] |
| BMI | 25.53 | 26.534 | 5.199 | [16.5,47.8] |

### PSQI data

For each participant, the total score and the individual component scores of the Pittsburgh sleep Quality Index (PSQI [54]) were collected from the most recent HCP data distribution. Due to extensive correlations across the seven components of the PSQI (see SOM2), prior to analyzing their relationship to brain structures, a data reduction was performed using a Principal Component Analysis (PCA; [63]) approach. The PCA was performed in JASP 2024 (v. 0.19.3; [64]), using an orthogonal varimax rotation and retaining all components with an eigenvalue greater than 1. This data reduction resulted in two orthogonal components explaining 50% of the variance. As shown in Tab. 1, the first principal component was loaded upon mainly by subjective sleep quality, sleep duration, and habitual sleep efficiency (with loadings equal to 0.62, 0.83, and 0.74, respectively). The second principal component was loaded upon by sleep onset latency, sleep disturbances, and use of sleep medications (with loadings equal to 0.67, 0.61, and 0.69, respectively). The daytime dysfunction PSQI component loaded weakly (*<* 0.36) on both both PCs. As is often the case with this measure, the PSQI total score, as well as the two PCs, are positively skewed (skewness coefficients: 1.029, 1.237, and 1.252, respectively) and not normally distributed, as assessed with a Shapiro-Wilk test of normality (W=0.931, p*<*0.001; W=0.907, p*<*0.001; and W=0.927, p*<*0.001, for the total score, PC1 and PC2, respectively). The ensuing exploration of associations between PSQI, the two principal components, demographic and biometric variables (i.e., sex, age, years of education, handedness, and BMI), and their interactions, are thus performed with a non-parametric approach (as described in the MRI data analysis section).

### MRI data

A total of 1,113 T1-weighted data were obtained from the HCP Young Adult repository. For each participant, the “T1w MPR1” dataset acquired as part of the “structural session” was employed. As described in the HCP Q3 release appendix (available at http://www.humanconnectome.org) these data were acquired with a Magnetization-Prepared Rapid Acquisition Gradient Echo (MPRAGE) sequence (TR = 2,400 ms, TE = 2.14 ms, TI = 1,000 ms, GRAPPA acceleration factor = 2). The acquisition resulted in an image size of 256*×*320*×*243 voxels (in the x, y, and z directions, respectively) and an isovoxel resolution of 0.7 mm.

### MRI data analysis

#### T1-weighted data preprocessing and segmentation

All MRI data analyses were performed with the FMRIB Software Library (FSL; [66]). T1-weighted data were first preprocessed using the `fsl_anat` script, which provides a general pipeline for processing anatomical images. For each T1-weighted image, the script reorients images to the standard (MNI) orientation (`fslreorient2std`), crops the image (`robustfov`), performs bias-field correction (RF/B1-inhomogeneity-correction; `FAST`; [67]), registration to standard space (linear and non-linear; `FLIRT` and `FNIRT`, [68, 69]), brain-extraction (`FNIRT`-based), tissue-type segmentation (`FAST`; [67]), and subcortical structure segmentation (`FIRST`; [57]). Once subcortical segmentations were completed, and before any analysis, the resulting subcortical meshes were individually inspected by the author to ensure correct structure segmentation (cf., Fig. 2). During this control, it emerged that segmentation for one participant failed, not producing any mesh output. That datapoint was thus excluded from any further analysis (leading to a final analyzed sample of 1,112 observations). Cortical segmentations were obtained by feeding the bias-corrected T1-weighted data to FSL Voxel Based Morphometry (VBM; [70]) scripts, an optimized VBM protocol[56]enclosed in the FSL distribution. Again, segmentations were visually inspected for accuracy prior to any analysis. All 1,112 T1-weighted images return satisfactory segmentations of the cortical gray matter (cf., Fig. 2).

#### T1-weighted data analysis

The single-smoothed subject cortical data (Gaussian smoothing with *σ* = 2, resulting in an approximate 4.71 mm full width half max [FWHM] Gaussian kernel) and subcortical meshes were entered into a group analysis. While both cortical and subcortical tissues are segmented in the subject’s native space, subcortical meshes are then co-registered to the standard MNI space for the purposes of group analysis. Conversely, cortical segmentations are co-registered to an *ad-hoc* sample-specific cortical template created, automatically, as an average of all the 1, 112 input data, for the purposes of performing the VBM group analysis. Each set of segmentations (i.e., cortical, subcortical) was entered into two group analyses. In the first analysis (henceforth, PSQI total score analysis), segmentations were regressed on the PSQI total score. In the second analysis (henceforth, PSQI PC analysis), segmentations were regressed on the two PC components described above. Each of these two statistical models also included a number of demographic (i.e., sex, age, handedness, years of education) and biometric (body mass index, BMI) sample characteristic variables. In addition, each analysis also included the interaction of each sample characteristic covarariate with the PSQI total score (in the PSQI total score analysis) and with PC1 and PC2 (in the PSQI PC analysis). Finally, each regression also included as an independent variable the subject’s total normalized brain volume (NBV; as calculated with `sienax`; [71]), in order to account for the effect of head size variability across individuals and to ensure that results reflect local shape change [65, 72].

Group analyses were performed with a non-parametric permutation testing approach [73] using `randomise` [74]. Significance was assessed against a criterion of *α ≤* 0.05 corrected for multiplicity with a family-wise cluster correction approach with Threshold Free Cluster Enhancement (`TFCE`; [75, 76]).

## Results

### PSQI scores

As shown in Fig. 1 and in Tab. 1, the PSQI total score in the analyzed sample had a mean of 4.8 and a median of 4. Using the conventional PSQI *>* 5 criterion [54], 31.8% of participants in the analyzed sample were classified as having poor sleep quality. The PSQI total score was not significantly different across sex (Mann-Whitney test [U_MW_]=147,234, p=0.246, effect size [rank-biserial correlation, r_rb_]=-0.04) and was not significantly correlated with age (Kendall’s Tau B [*τ*_*B*_]=-0.019, p=0.379, effect size [Fischer’s z]=-0.019). However, PSQI was significantly associated with handedness (*τ*_*B*_=-0.064, p=0.003, Fisher’s z=-0.064), with greater right-handed lateralization showing smaller PSQI total scores (i.e., better sleep quality), years of education (*τ*_*B*_=-0.108, p*<*0.001, Fisher’s z=-0.108), with more years of education being associated with better sleep quality, and BMI (*τ*_*B*_=0.072, p*<*0.001, Fisher’s z=0.072), with greater biometric values being associated with worse sleep quality (see Fig. SOM1 for a depiction of the corrrelations between PSQI total score, PSQI PC1, PC2, and all sample characteristic covariates).

**Fig. 1.**
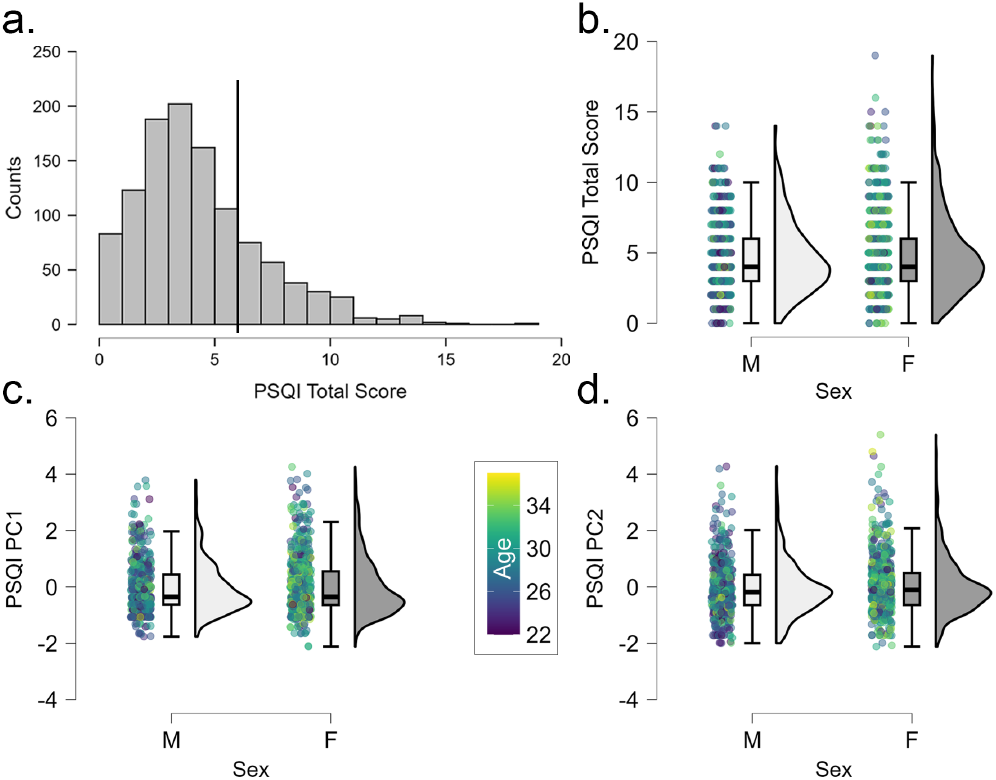
PSQI data. (a) Distribution of PSQI total scores in the analyzed sample. The vertical black line indicates the conventional criterion discriminating “poor sleepers” (i.e., PSQI total score *>* 5) from “good sleepers” (i.e., PSQI total score *≤* 5; [54]). (b-d) Distribution of PSQI total score, PSQI PCA PC1, and PC2 by sex and age group. (No significant differences were observed in the distribution of PSQI total score, or either PCA component, across sex or age group.)

**Fig. 2.**
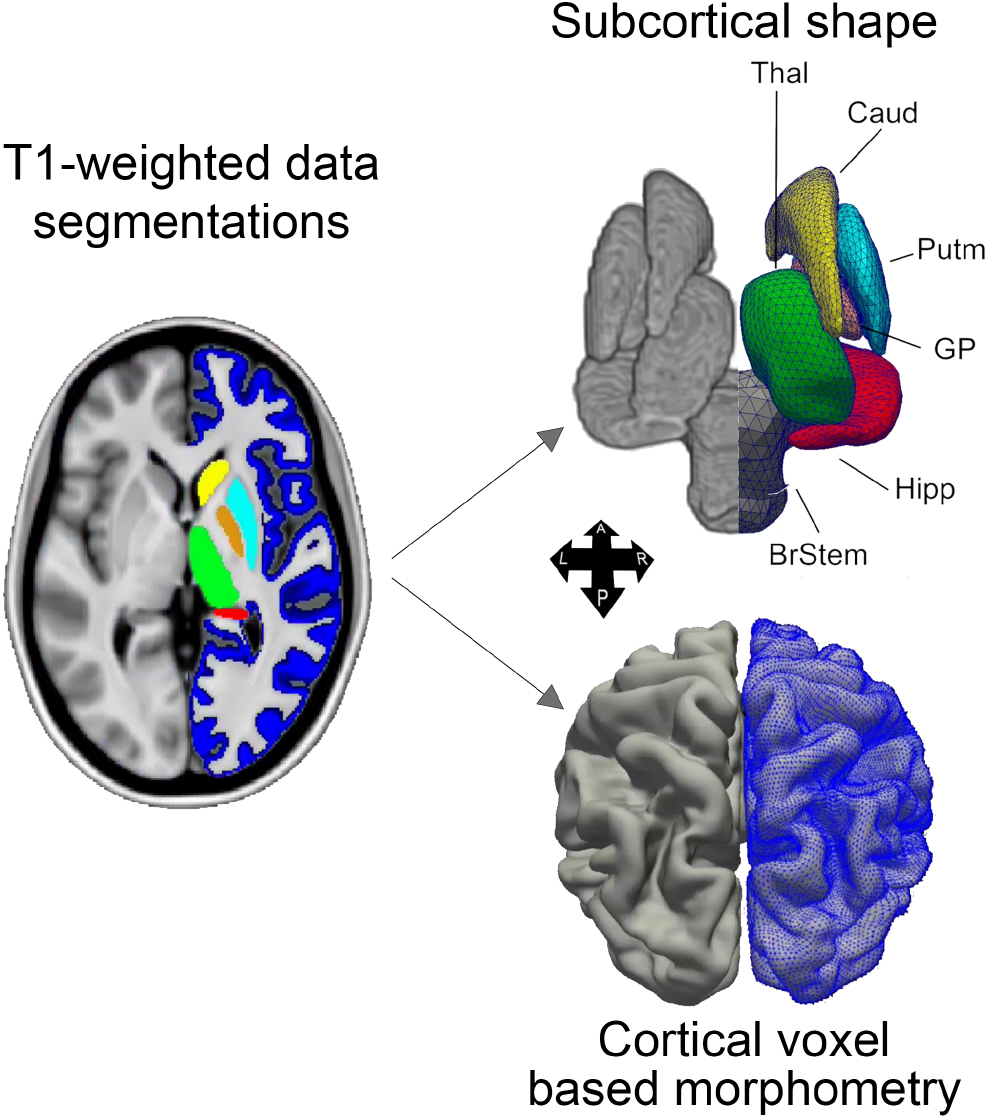
Methods. Left: Illustration of cortical and subcortical segmentations of the T1-weighted data. Right: Illustration of 3D subcortical (top) and cortical (bottom) reconstructed meshes. (Image partially adapted from [65])

With respect to the two PSQI PCs, in terms of their interpretation, both scores were positively associated with higher PSQI scores, and therefore worse sleep quality (*τ*_*B*_=0.584, p*<*0.001, Fisher’s z=0.669 and *τ*_*B*_=0.478, p*<*0.001, Fisher’s z=0.521, for PC1 and PC2, respectively). Similarly to the PSQI total score, neither PC was significantly different across sex (U_MW_=152,589, p=0.884, r_rb_=-0.005 and U_MW_=146,676, p=0.209, r_rb_=-0.044, for PC1 and PC2, respectively) and neither was significantly correlated with age (*τ*_*B*_=-0.001, p=0.953, Fischer’s z=-0.001 and *τ*_*B*_=-0.008, p=0.689, Fischer’s z=-0.008, for PC1 and PC2, respectively). Interestingly, PC1’s associations with the remaining covariate measures paralleled those reported for the PSQI total score (i.e., handedness: *τ*_*B*_=-0.058, p=0.006, Fischer’s z=-0.058; BMI: *τ*_*B*_=0.103, p*<*0.001, Fischer’s z=0.104; education: *τ*_*B*_=0.103, p*<*0.001, Fischer’s z=0.103). In contrast, no significant associations were observed between PC2 and any of the remaining covariates (i.e., handedness: *τ*_*B*_=-0.012, p=0.566, Fischer’s z=-0.012; BMI: *τ*_*B*_=0.003, p=0.884, Fischer’s z=0.003; education: *τ*_*B*_=-0.023, p=0.303, Fischer’s z=-0.023). (See Fig. SOM1 for a graphical depiction of the PCs correlations with PSQI total score and covariates.)

### MRI results

As shown in Fig. 3, the PSQI total score was associated, inversely, with subcortical shape in portions of the bilateral caudate and putamen, implying an association between poorer sleep and smaller shapes in these regions (cf., Tab. SOM3 for precise neuroanatomical localization of significant clusters). No positive associations were detected between the PSQI total score and subcortical shape. No significant associations were detected, either positive or negative, between voxelwise cortical morphometry and the PSQI total score. Mirroring the correlation observed for the PSQI total score, as shown in Fig. 3, the PSQI PC1 was also negatively associated with subcortical shape in similar portions of the bilateral caudate and putamen, as well as additional portions of the left pallidum (see Tab. SOM4 for precise neuroanatomical localization of significant clusters), again implying an association between smaller subcortical shapes and poorer sleep quality, particularly with respect to subjective sleep quality, sleep duration, and habitual sleep efficiency. No positive associations were detected between PC1 and subcortical shape. As for the PSQI total score, no significant associations were detected, either positive or negative, with cortical morphometry. Finally, PC2, the PSQI principal component which was loaded upon maximally by sleep onset latency, sleep disturbance, and use of sleep medications, did not exhibit any significant association, either positive or negative, with either subcortical shape or cortical morphometry.

**Fig. 3.**
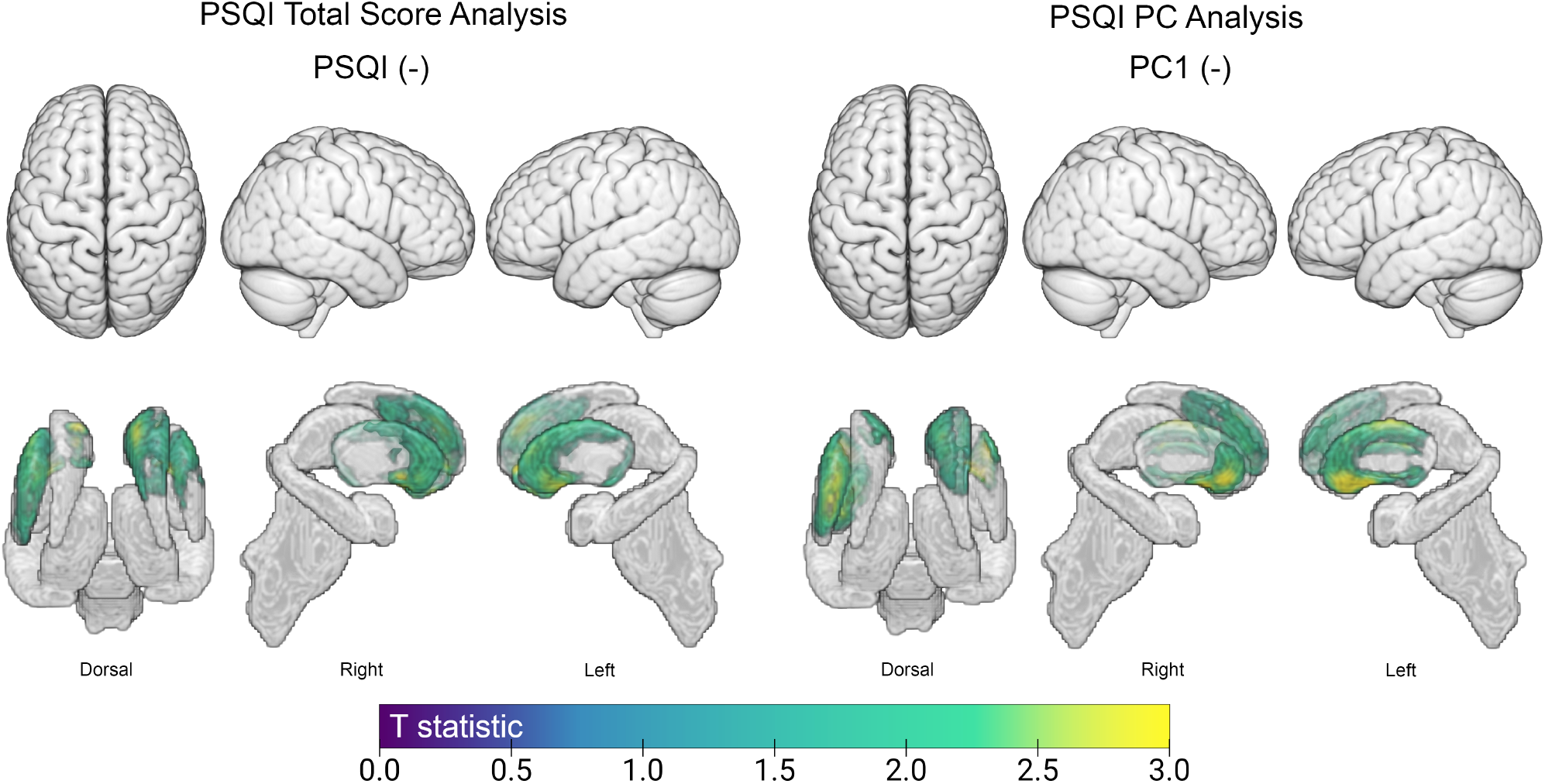
Correlates of sleep quality measures. Depiction of all voxels significantly associated with PSQI total score regressor (left), in the PSQI total score analysis, and with the PSQI PC1 regressor (right), in the PSQI PC analysis. As shown, only negative associations were detected between subcortical shape and the two sleep quality variables. No significant associations were detected in cortex for any of the sleep quality measures, neither positive nor negative, and no significant associations were observed for the PC2 component, in either cortex or subcortical regions, either positive or negative. (Note: colored areas imply a significant association, corrected for multiplicity; gray areas imply no significant associations.) See Tabs. SOM3 and SOM4 for detailed localization of each area of significant association.

In addition to the associations reported above for the regressors of interest (i.e., PSQI total score and the PSQI PCs), significant cortical and subcortical associations were also detected when assessing the interaction of each regressors of interest with demographic and biometric sample characteristics. As shown in Fig. 4, significant associations were uncovered between subcortical shape and the interaction of the PSQI total score with BMI, education, and handedness (see also Tab. SOM3). Specifically, a negative relationship was observed between the interaction of PSQI and BMI and shape in portions of the bilateral pallidum and hippocampus, as well as between the interaction of PSQI and handedness and portions of the bilateral caudate, hippocampus, and right amygdala. Conversely, a positive association was observed between the interaction of PSQI and years of education, and portions of the bilateral caudate, right thalamus and right pallidum.

**Fig. 4.**
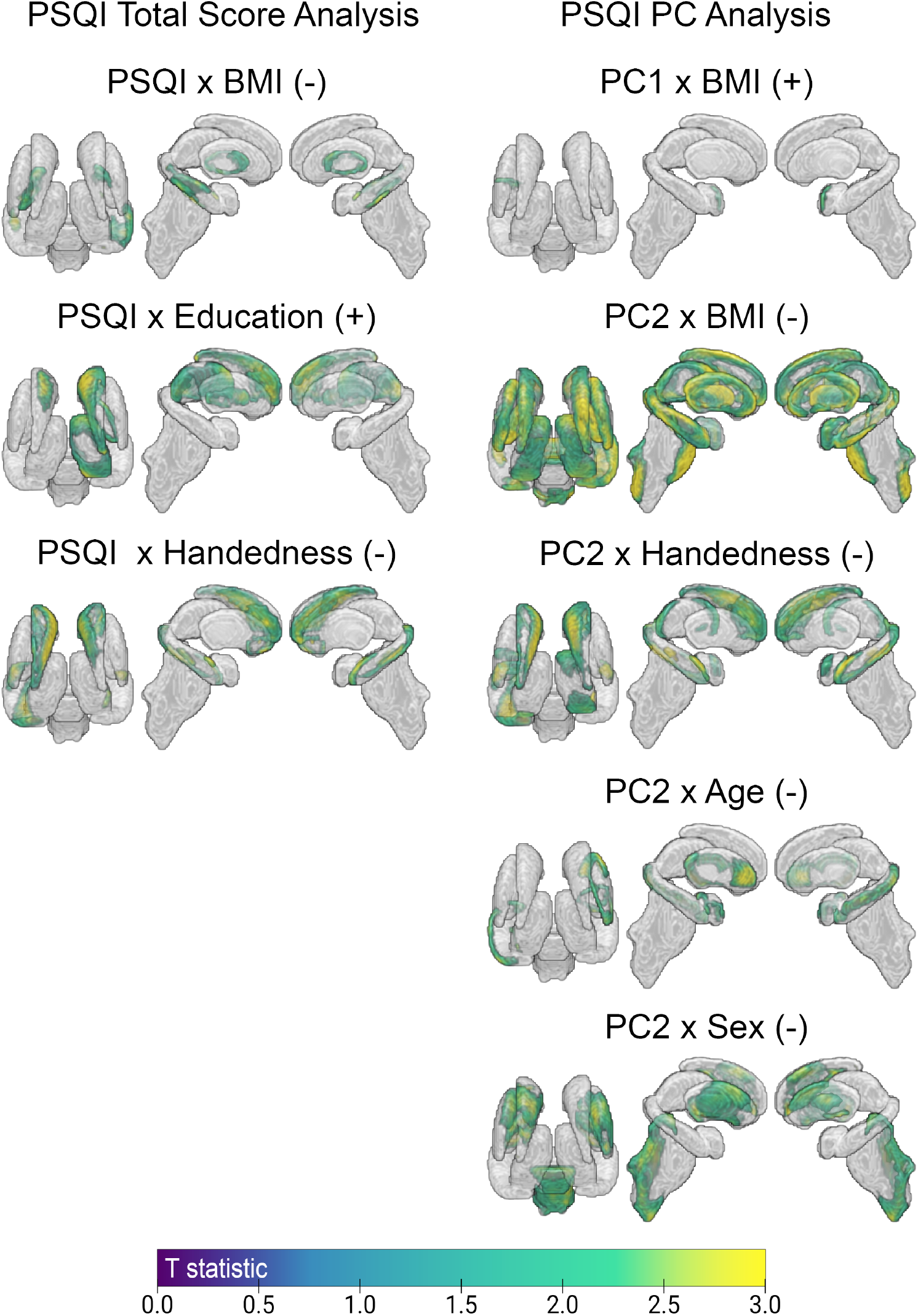
Subcortical correlates of interactions with sample characteristic variables. Left: Depiction of all voxels significantly associated with the interaction of PSQI total score with each demographic and biometric covariate (i.e., sample characteristics variables). Right: Depiction of all voxels significantly associated with the interaction of PC1 and PC2, respectively, with each demographic and biometric covariate. See Tab. SOM3 and SOM4, respectively, for anatomical localization and statistics of each cluster. (Areas shown in color indicate significant voxels after correction for multiple comparisons. Areas shown in gray imply no significant result. ‘+’ indicates to a positive association; ‘-’ indicates a negative association.)

Significant associations with interaction terms were also uncovered in the PSQI PC analysis (cf., Fig. 4 and Tab. SOM4). Specifically, the interaction of PC1 and BMI was positively correlated with subcortical shape in a small portion of the left amygdala. PC2, despite showing no direct association with any of the sleep quality regressors (i.e., PSQI total score, PSQI PCs), did exhibit multiple significant associations with the interaction of the sleep quality regressors with age, BMI, handedness, and sex. Specifically, the interaction of PC2 and age associated negatively with subcortical shape in portions of the left hippocampus, right caudate and amygdala. The interaction of PC2 and BMI revealed extensive negative associations with subcortical shape in large portions of the bilateral thalamus, caudate, putamen, pallidum, hippocampus, as well as the left amygdala and the brainstem. The interaction of PC2 and handedness also revealed a negative association with shape across bilateral caudate, hippocampus, right thalamus, and left amygdala. Similarly, the interaction of PC2 and sex was also associated with subcortical shape in broad portions of the bilateral pallidum, left caudate and putamen, as well as the brainstem. Finally, a small positive association was detected between subcortical shape in the left putamen and the interaction of PC2 and education.

As shown in Fig. 5, despite the absence of any significant association between cortical voxel based morephometry and the main sleep quality regressors (i.e., PSQI total score and PSQI PCs), a number of significant associations were detected between cortical morphometry and PC interaction terms. Specifically, the interaction of PC1 and age revealed positive associations in the bilateral insula, the left parahippocampal gyrus and temporal pole, as well as medial occipital cortex, in the occipito-parietal sulcus and lingual gyrus. In addition, two small negative associations were found, between PC1 and age, in left posterior parietal cortex. The interaction of PC2 and education was negatively associated with with morphometry measures in the right anterior insular cortex, while the interaction of PC2 and sex was associated with morphometry measures in two foci in the medial wall of frontal cortex, in the regions of the supplementary motor area and the paracingulate gyrus. (See Tab. SOM5 for precise neuroanatomical localization of significant clusters.)

**Fig. 5.**
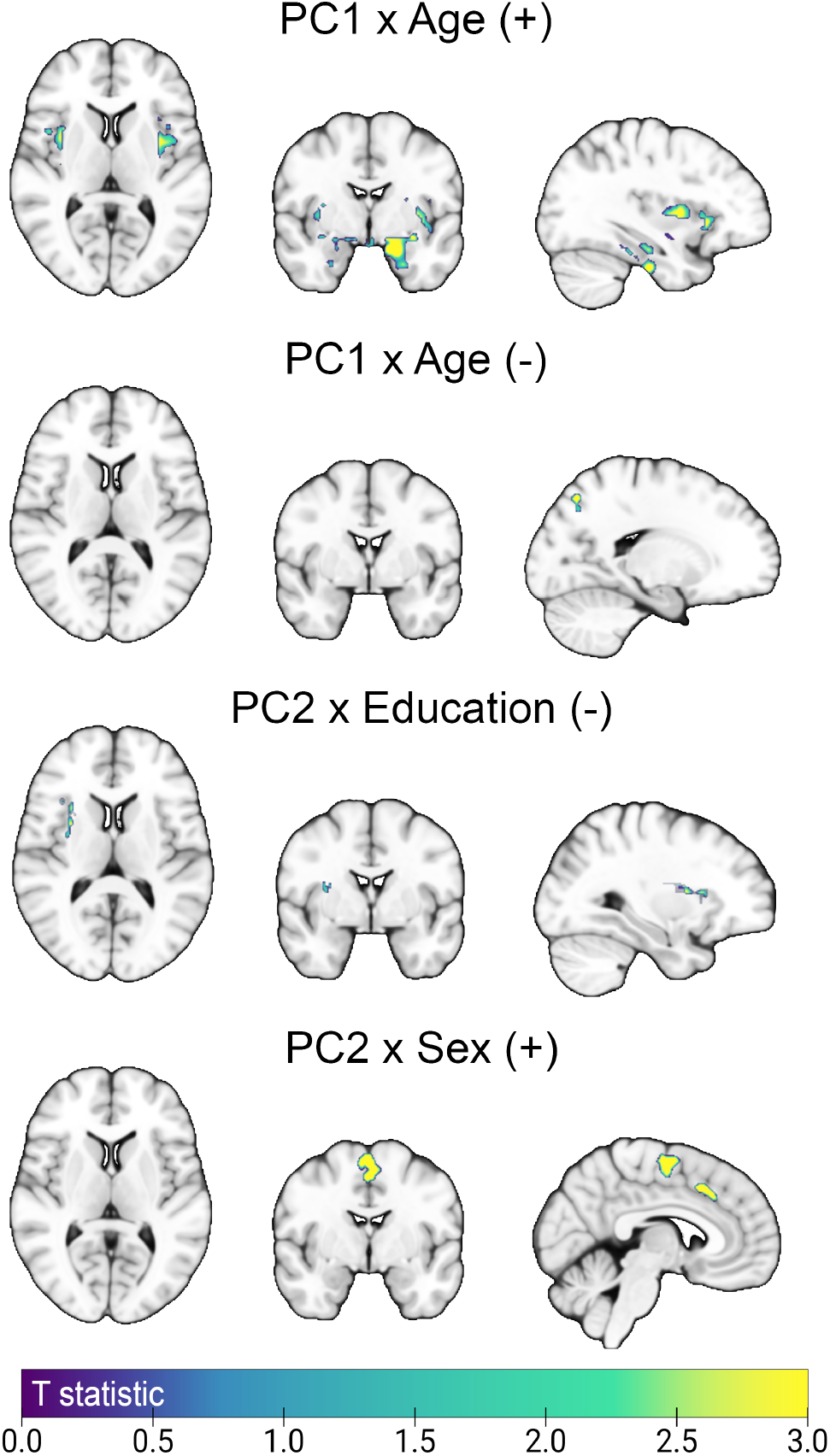
Cortical correlates of interactions with sample characteristic variables. Depiction of all voxels significantly associated with the interaction of PSQI PC1 and PC2 with each demographic and biometric covariate (i.e., sample characteristics variables). (Note that no significant associations were found between cortical morphometry and the interaction of PSQI total score and sample characteristic variables.) See Tab. SOM5 for anatomical localization and statistics of each significant cluster. (Areas shown in color indicate significant voxels after correction for multiple comparisons. Areas shown in gray imply no significant result. ‘+’ indicates to a positive association; ‘-’ indicates a negative association.)

## Discussion

The present work shows that lower self-reported sleep quality and, in particular, subjective sleep quality, sleep duration, and habitual sleep efficiency, is associated with smaller subcortical shapes in the bilateral striatum and the left pallidum, after parceling out any variance explained by sample characteristics (i.e., demographic and biometric covariates) and their interaction with sleep quality measures.

Long known to serve a key role in the control of motor behavior [77], the contributions of the basal ganglia to non-motor function have only come to the forefront relatively recently [52, 78, 79], together with a renewed understanding of the complex circuitry tying these regions together and within broader cortical-subcortical circuits [80, 81]. Indeed, clinical data show that patients with striatal dysfunction syndromes, such as Parkinson’s and Huntington Disease, exhibit high rates of sleep disturbances [17–20, 82], something that is likely to be mediated, at least in part, by dopamine mechanisms [83]. Patients suffering from insomnia disorders show an association between shape change in many of these regions and sleep quality measures [60]. Interestingly, the availability of striatal dopamine in D2/D3 receptors has been associated with sleep quality in healthy adults [84]. Recent work in a large cohort of healthy volunteers also highlighted the association between neural tissue density in these regions (as assessed with diffusion tensor imaging) and sleep duration [85]. Consistent with these results, and the present findings, lesion studies in the rodent model have documented alteration of sleep-wake behavior following striatal damage [86, 87]. In particular, striatal lesions have been shown to lead to greater sleep-wake behavior fragmentation, including greater state transitions, loss of ultra-long wake bouts, and diminished diurnal sleep-wake variation [88]. It is not surprising, then, that despite significant heterogeneity in findings and measures (e.g., PSQI versus non-normative sleep duration), neuroanatomical variability in these regions has been previously associated to measures of sleep quality in healthy individuals [48–50]. Also conventionally conceived as a relay in the control of motor behavior [89– 91], a growing amount of evidence links the pallidum – and its external portion in particular – to the regulation of sleep-wake behavior and electrophysiological arousal [78, 79], likely *via* dopaminergic neuromodulation [92]. Indeed, as found with the striatum, excitotoxic lesions of the external portion of the pallidum, in rodent models, lead to increased sleep fragmentation, with increased sleep-wake transitions and decreased sleep bouts [88, 93]. Conversely, optogenetic and electical neuromodulation of this region results in a broad increase of time spent sleeping [93, 94]. Consistent with these findings, in humans, the pallidum has been associated with behavioral and/or electrocortical arousal in healthy volunteers undergoing general anesthesia [95], and in severe brain injury patients [65, 72]. Intriguingly, and mirroring the rodent work, deep brain stimulation of the external portion of the pallidum in a single parkinsoninan patient dramatically reduced their sleep-related symptomatology [96].

The interaction of sleep quality, demographic, and biometric factors highlighted several additional associations with brain neuroanatomy in both subcortical (i.e., hippocampus, thalamus, amygdala, and brainstem) and cortical regions (i.e., sublobar, insular, and mesial cortices). Many of these areas have been previously associated with aspects of sleep quality. Hippocampal morphometry, for example, has been previously shown to have a negative relation with sleep quality indices, both cross-sectionally [49, 50] and longitudinally [40]. Similarly, prior work using diffusion tensor imaging also highlighted an association between (right) hippocampus and greater tissue density (as measured with mean diffusivity) [85]. Thalamus, and its interactions with cortex, have long been implicated in sleep-wake transitions, as a key target of ascending arousal system [97, 98]. Large sample population studies have revealed associations between non-normative sleep duration and macrostructural and microstructural brain outcomes, including thalamus (as well as hippocampus, putamen, and accumbens) [48, 49]. Sleep disturbances have also been associated with thalamic abnormalities and/or dysfunction in healthy volunteers [99] and across many mental health and neuropsychiatric disorders [49, 100–102]. Furthermore, thalamic stroke and infraction are known to lead to sleep impairments [103, 104] and long-term sleep deprivation has been associated with decreased thalamic volume [105]. Importantly, in patients with insomnia, clinical features of the pathology are also associated with the connectivity between thalamus and many of the regions uncovered in the present work including cingualte cortex, hippocampus, and striatum [100]. Atrophy and connectivity dysfunction in the amygdala has also been associated with sleep disturbances across a number of conditions [50, 106–109], dissociably from its co-morbid association with anxiety [110]. Finally, structural and functional aspects of insular cortex, including its interactions with cingulate cortex [111], have been previously shown to be disrupted by sleep deprivation [112] as well as sleep disorders [113–116].

Despite this superficial consistency with prior work, it cannot be ignored that in the present work these associations are specific to the interaction of sleep quality measures and sample characteristics (i.e., demographic and biometric variables). This raises an important methodological issue in addition to those previously discussed [51]: while almost all studies make use of a number – often large – of covariates in assessing their effects of interest, many do not assess any interaction [e.g., 38, 49, 85, 99, 117] and, those that do, only include one or two interactions of interest (e.g., sex, age) [e.g., 34, 40, 48, 118]. In the context of a regression-type analysis, this can mean that variance might be misallocated to some extent, potentially resulting in spurious results. One interesting case in point is the previously reported association between the PSQI*×*Sex interaction and local neuroanatomical features in frontal cortex, along with sex-specific differences in the parahippocampus and parietal cortices [34]. In the present analysis no such effect was observed. Yet, as shown in Fig. 5, a subset of these regions were observed to associate – albeit with important lateralization differences – with the interaction of PSQI and age. Of course, whether the discrepancy is due to the use of different statistical models (e.g., inclusion of different interaction terms) or to other factors remains to be understood. Nonetheless, it is noteworthy that, while the PSQI total score by sex interaction did not exhibit any significant association with either cortical or subcortical morphometry in the analysis presented above (cf., Figs. 4,5 and Tabs. SOM3-SOM5), when the analysis was re-run omitting all interaction terms with the exception of one interaction of interest (i.e., PSQI*×*Sex), significant associations were detected in multiple subcortical regions (see Fig. SOM2 for a depiction of the result). This finding further highlights the multifaceted nature of brain-sleep quality relations as well as the need for considering more sophisticated approaches, in testing such associations, that can better model complex structures (e.g., structural equation modeling [119, 120], causal discovery analysis [121], Bayesian network analysis [122]).

Although not an aim of the present work, the results reported above also contribute to the discussion concerning whether the factor structure of the PSQI includes more than one dimension and, if it does, how individual items/components of the instrument distribute across dimensions [123–125]. Here, an exploratory PCA with varimax rotation was employed as a statistical device for the sole purposes of obtaining (rotated) components which, if more than one, would be uncorrelated with each other, thus avoiding the issue of excessive multicollinearity in the general linear model analysis [126]. Nonetheless, it is interesting to observe that the two components derived from this approach exhibited a different correlational structure with demographic and biometric variables, brain measures, and their interactions. PC1, which was loaded maximally upon by subjective sleep quality, sleep duration, and habitual sleep efficiency, exhibited a similar correlational structure with demographic and biometric covariates to that of the PSQI total score. Both were similarly and significantly associated with handedness, BMI, and years of education. Conversely, PC2, which was loaded upon maximally by sleep onset latency, sleep disturbance, and use of sleep medications, exhibited no such associations (cf., Tab. 1 and Fig. SOM1). The two PCs also exhibited a different correlational structure with brain measures (cf., Fig. 4,5 and Tab. SOM3-SOM5). Indeed, PC1 exhibited a similar association with subcortical areas as that observed for the PSQI total score. Conversely, PC2 did not show any significant associations with cortical or subcortical morphometry. It is also relevant that while the PSQI total score failed to show any significant association with cortical morphometry, both in the context of the main PSQI total score regressor as well as its interaction with sample characteristics, PC1 and PC2 did return significant cortical morphometry associations with select interactions, indicating that they are capturing different aspects of the relationship between neuroanatomical variability and self-reported sleep quality measures. It should be highlighted, however, that all the above considerations result from data-driven observations, not systematic assessments of the dimensionality of the PSQI, and should thus be interpreted as such.

Finally, in assessing the present findings, one must take stock of a number of limitations tied to the approach employed. First, the segmentation technique employed here[57] cannot distinguish a number of regions and subregions (e.g., parabrachial nucleus [127, 128], internal versus external segments of the globus pallidus [129–131], reticular [132– 134], anterior and mediodorsal [135] nuclei), cell populations (e.g., cell versus matrix thalamic neurons [136]), as well as neurotransmetter-specific (e.g., glutamatergic, GABAergic [137]) neuromodulatory systems, that are believed to play an important role in the regulation of sleep-wake behavior. Second, the morphometric approach employed centers on local cortical and subcortical structures, but is blind to other factors (e.g., white matter integrity, glymphatic system, perivascular space), which have been proposed to play a role in sleep [117, 138–141]. Third, the present work is focused entirely on neuroanatomical variability, yet there exist known associations between changes in brain function and sleep quality [58, 142– 147]. It will be interesting, in future work, to delineate the degree to which (if any) the known associations between brain network function and perceived sleep quality [e.g., 58, 148] is related to the neuroanatomical variability observed here. Finally, while longitudinal studies have reported atrophy and smaller volumes over time to be associated with decreasing sleep quality [e.g., 36, 40, 118], the present findings are cross-sectional in nature and do not permit claims of causality in the uncovered brain-sleep quality associations.

## Conclusion

This study demonstrates a robust association between conventional self-reported measures of sleep quality [54] and subcortical morphometry, particularly in the bilateral caudate, putamen, and left pallidum. These findings provide evidence for the importance of the basal ganglia in sleep regulation in healthy, young, individuals, reinforcing prior work linking striatal function to sleep disturbances in both healthy and clinical populations [52, 78, 79]. While cortical morphometry did not directly correlate with sleep quality measures, interactions with demographic and biometric factors were observed, emphasizing the multifaceted nature of brain-sleep relationships. These results underscore the importance of considering both direct and interaction effects when studying neuroanatomical correlates of sleep. Finally, in light of the rising availability of non-invasive deep brain neuromodulatory techniques (e.g., low-intensity transcranial focused ultrasound [149–151]), the potential identification of subcortical structures as sleep-related hubs could be investigated causally and, should such a link be established, could present an opportunity for targeted interventions aimed at improving sleep quality in both healthy individuals and clinical populations [52, 96].

## Data availability statement

The data underlying this article are publicly available at https://www.humanconnectome.org/study/hcp-young-adult. This manuscript is made available on BiorXiv as preprinthttps://doi.org/10.1101/2024.05.29.596530.

## Acknowledgments

The author would like to acknowledge the UCLA Institute for Digital Research and Education (IDRE) for making temporarily available to investigators large computational space and resources for allowing processing and analysis of large datasets.

## Author contributions statement

M.M.M. conceived the experiment, analysed the data, and wrote the manuscript.

## Supplementary Online Materials

**Table SOM1.** Sample demographic and PSQI information broken down by sex. (Abbreviations: Med., median; sd., standard deviation; PC1, PSQI Principal Component 1; PC2, PSQI Principal Component 2; subj. sleep qual., subjective sleep quality; sleep dur., sleep duration; sleep eff., sleep efficiency; sleep ons. lat., sleep onset latency; sleep disturb., sleep disturbances; sleep med., use of sleep medications; daytime dysf., daytime dysfunction; U_MW_, Mann-Whitney U test for independent samples; r_rb_, rank-biserial correlation. Significance level of the Mann-Whitney test between female and male: *** indicates p*<*0.001; ** indicates p*<*0.01; * indicates p*<*0.05. See Tab. 1 for sample-wide statistics.)

| Variables | Female (N=605) | | | Male (N=507) | | | $U_{MW}$ | $r_{rb}$ |
| --- | --- | --- | --- | --- | --- | --- | --- | --- |
|  | Med | Mean | sd | Med | Mean | sd |  |  |
| PSQI Total Score | 4 | 4.942 | 2.933 | 4 | 4.639 | 2.528 | 147,234 | -0.04 |
| Subj. sleep qual. (PC1) | 1 | 0.899 | 0.66 | 1 | 0.888 | 0.604 | 153,910 | 0.004 |
| Sleep dur. (PC1) | 0 | 0.559 | 0.817 | 0 | 0.582 | 0.814 | 156,488 | 0.02 |
| Sleep eff. (PC1) | 0 | 0.534 | 0.846 | 0 | 0.363 | 0.719 | 137209*** | -0.105 |
| Sleep ons. lat. (PC2) | 1 | 0.975 | 0.833 | 1 | 0.957 | 0.799 | 152,612 | -0.005 |
| Sleep disturb. (PC2) | 1 | 1.129 | 0.492 | 1 | 1.049 | 0.474 | 143149** | -0.067 |
| Sleep med. (PC2) | 0 | 0.261 | 0.717 | 0 | 0.209 | 0.625 | 149,911 | -0.023 |
| Daytime dysf. (-) | 1 | 0.585 | 0.642 | 1 | 0.592 | 0.626 | 154,947 | 0.01 |
| Age | 30 | 29.55 | 3.6 | 28 | 27.901 | 3.611 | 114060*** | -0.256 |
| Handedness | 80 | 70.388 | 43.761 | 75 | 60.424 | 44.17 | 116345*** | -0.241 |
| Education | 16 | 15.015 | 1.815 | 16 | 14.822 | 1.776 | 140547* | -0.08 |
| BMI | 24.92 | 26.279 | 5.785 | 26.11 | 26.836 | 4.388 | 173375*** | 0.132 |

**Table SOM2.**
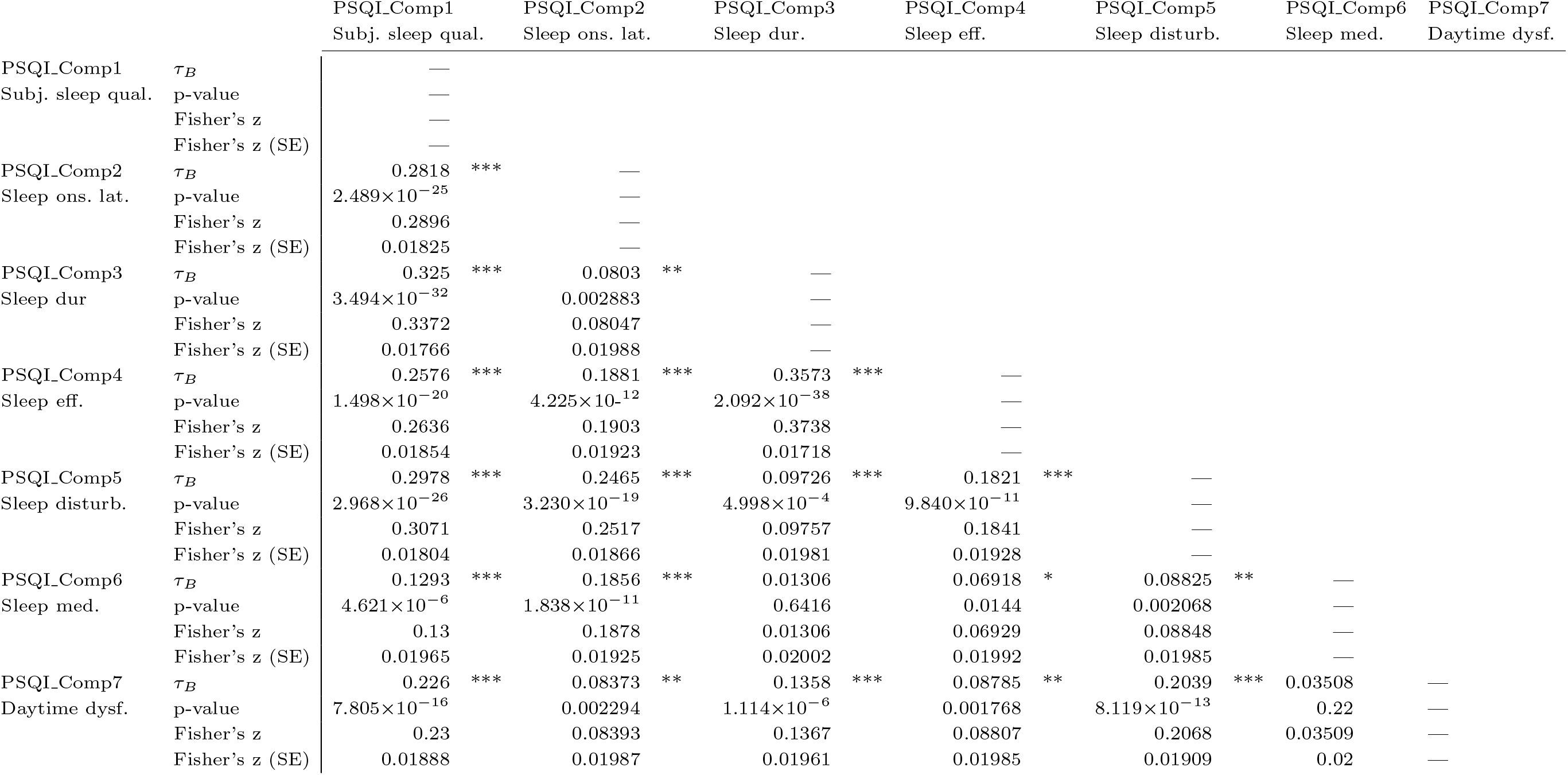
Correlations between the PSQI components. (Abbreviations: *τ*_*B*_, Kendall’s Tau B; Fisher’s z (SE), standard error of the Fisher’s z effect size. All other abbreviations are as in SOM1.)

**Table SOM3.**
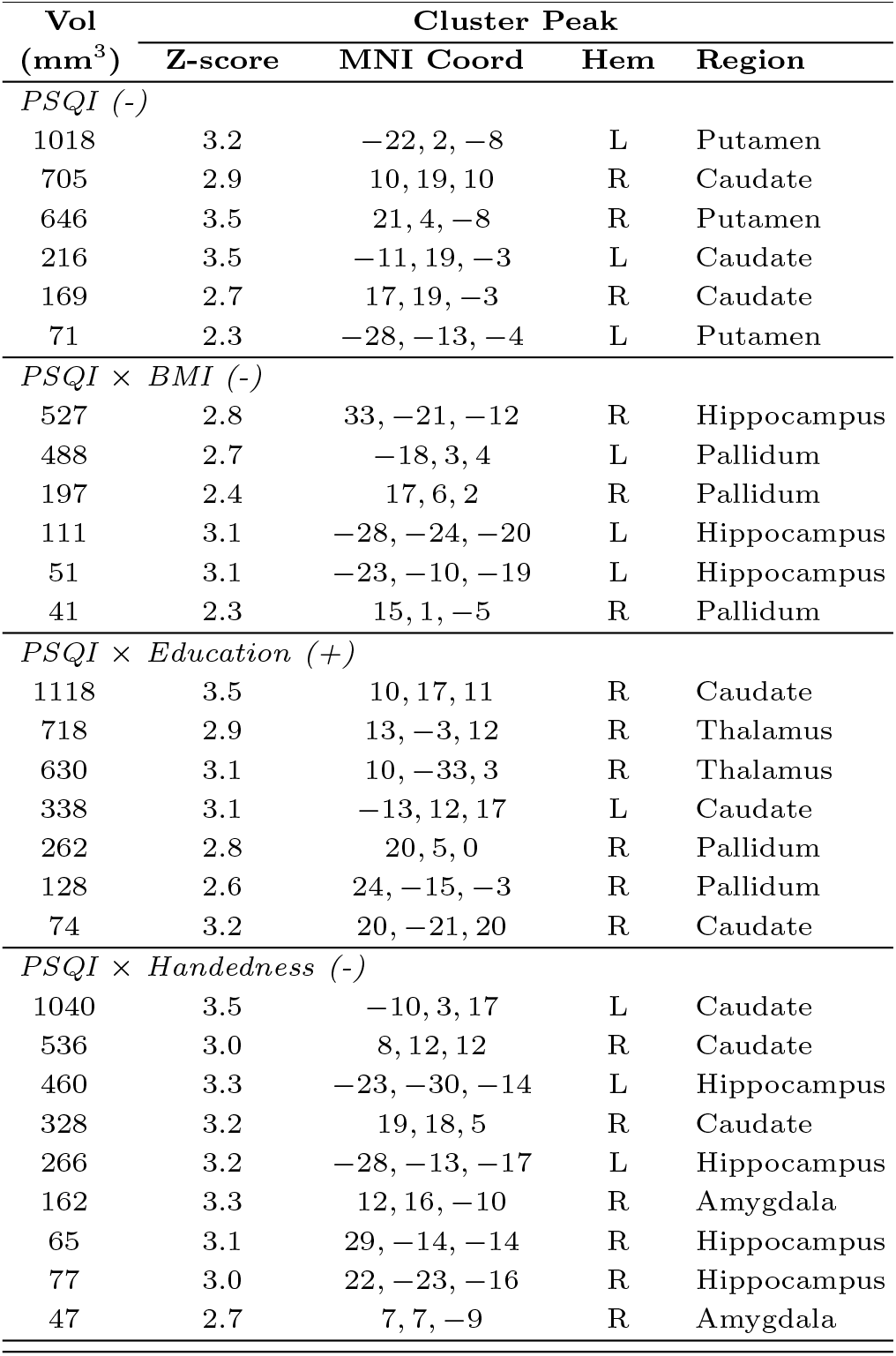
Subcortical Results. Significant associations between variables included in the PSQI total score analysis and subcortical shape. See Fig. 4. For each cluster, volume, peak Z-score and MNI coordinates, hemisphere, and anatomical label are shown. (Abbreviations and symbols: Vol, volume; Coord, coordinates; Hem, hemisphere; L, left; R, right; ‘-’ represents a negative association; ‘+’ represents a positive association.)

| Vol<br>(mm<sup>3</sup>) | Z-score | MNI Coord | Hem | Region |
| --- | --- | --- | --- | --- |
| *PSQI (-)* |  |  |  |  |
| 1018 | 3.2 | -22, 2, -8 | L | Putamen |
| 705 | 2.9 | 10, 19, 10 | R | Caudate |
| 646 | 3.5 | 21, 4, -8 | R | Putamen |
| 216 | 3.5 | -11, 19, -3 | L | Caudate |
| 169 | 2.7 | 17, 19, -3 | R | Caudate |
| 71 | 2.3 | -28, -13, -4 | L | Putamen |
| *PSQI × BMI (-)* |  |  |  |  |
| 527 | 2.8 | 33, -21, -12 | R | Hippocampus |
| 488 | 2.7 | -18, 3, 4 | L | Pallidum |
| 197 | 2.4 | 17, 6, 2 | R | Pallidum |
| 111 | 3.1 | -28, -24, -20 | L | Hippocampus |
| 51 | 3.1 | -23, -10, -19 | L | Hippocampus |
| 41 | 2.3 | 15, 1, -5 | R | Pallidum |
| *PSQI × Education (+)* |  |  |  |  |
| 1118 | 3.5 | 10, 17, 11 | R | Caudate |
| 718 | 2.9 | 13, -3, 12 | R | Thalamus |
| 630 | 3.1 | 10, -33, 3 | R | Thalamus |
| 338 | 3.1 | -13, 12, 17 | L | Caudate |
| 262 | 2.8 | 20, 5, 0 | R | Pallidum |
| 128 | 2.6 | 24, -15, -3 | R | Pallidum |
| 74 | 3.2 | 20, -21, 20 | R | Caudate |
| *PSQI × Handedness (-)* |  |  |  |  |
| 1040 | 3.5 | -10, 3, 17 | L | Caudate |
| 536 | 3.0 | 8, 12, 12 | R | Caudate |
| 460 | 3.3 | -23, -30, -14 | L | Hippocampus |
| 328 | 3.2 | 19, 18, 5 | R | Caudate |
| 266 | 3.2 | -28, -13, -17 | L | Hippocampus |
| 162 | 3.3 | 12, 16, -10 | R | Amygdala |
| 65 | 3.1 | 29, -14, -14 | R | Hippocampus |
| 77 | 3.0 | 22, -23, -16 | R | Hippocampus |
| 47 | 2.7 | 7, 7, -9 | R | Amygdala |

**Table SOM4.**
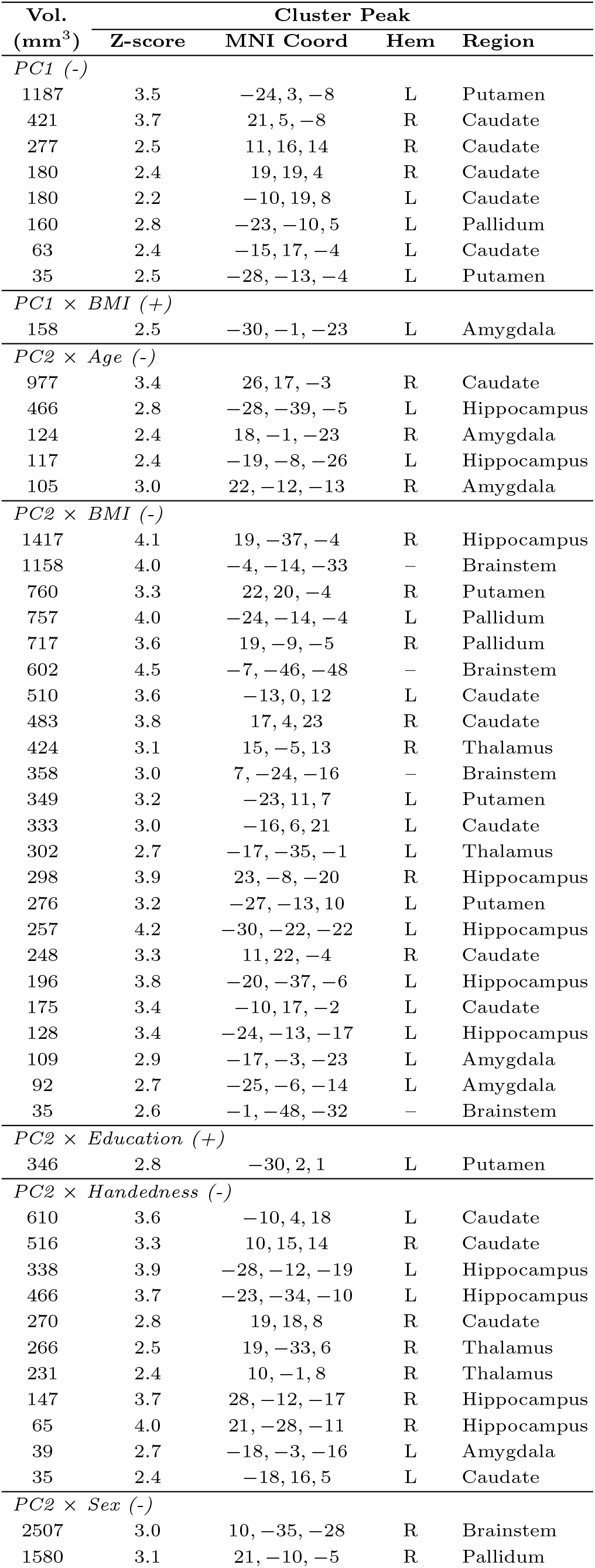
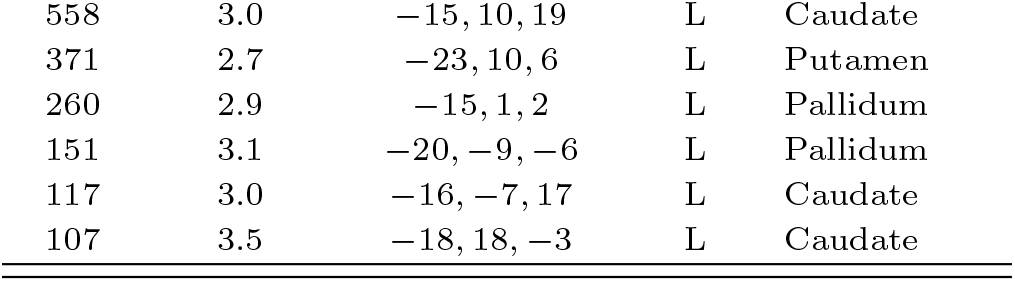
Subcortical Results. Significant associations between variables included in the PSQI PC analysis and subcortical shape. See Fig. 4. (Abbreviations are the same as in Tab. SOM3.)

| Vol.<br>(mm<sup>3</sup>) | Cluster Peak |  |  |  |
| --- | --- | --- | --- | --- |
|  | Z-score | MNI Coord | Hem | Region |
| *PC1 (-)* |  |  |  |  |
| 1187 | 3.5 | -24, 3, -8 | L | Putamen |
| 421 | 3.7 | 21, 5, -8 | R | Caudate |
| 277 | 2.5 | 11, 16, 14 | R | Caudate |
| 180 | 2.4 | 19, 19, 4 | R | Caudate |
| 180 | 2.2 | -10, 19, 8 | L | Caudate |
| 160 | 2.8 | -23, -10, 5 | L | Pallidum |
| 63 | 2.4 | -15, 17, -4 | L | Caudate |
| 35 | 2.5 | -28, -13, -4 | L | Putamen |
| *PC1 × BMI (+)* |  |  |  |  |
| 158 | 2.5 | -30, -1, -23 | L | Amygdala |
| *PC2 × Age (-)* |  |  |  |  |
| 977 | 3.4 | 26, 17, -3 | R | Caudate |
| 466 | 2.8 | -28, -39, -5 | L | Hippocampus |
| 124 | 2.4 | 18, -1, -23 | R | Amygdala |
| 117 | 2.4 | -19, -8, -26 | L | Hippocampus |
| 105 | 3.0 | 22, -12, -13 | R | Amygdala |
| *PC2 × BMI (-)* |  |  |  |  |
| 1417 | 4.1 | 19, -37, -4 | R | Hippocampus |
| 1158 | 4.0 | -4, -14, -33 | - | Brainstem |
| 760 | 3.3 | 22, 20, -4 | R | Putamen |
| 757 | 4.0 | -24, -14, -4 | L | Pallidum |
| 717 | 3.6 | 19, -9, -5 | R | Pallidum |
| 602 | 4.5 | -7, -46, -48 | - | Brainstem |
| 510 | 3.6 | -13, 0, 12 | L | Caudate |
| 483 | 3.8 | 17, 4, 23 | R | Caudate |
| 424 | 3.1 | 15, -5, 13 | R | Thalamus |
| 358 | 3.0 | 7, -24, -16 | - | Brainstem |
| 349 | 3.2 | -23, 11, 7 | L | Putamen |
| 333 | 3.0 | -16, 6, 21 | L | Caudate |
| 302 | 2.7 | -17, -35, -1 | L | Thalamus |
| 298 | 3.9 | 23, -8, -20 | R | Hippocampus |
| 276 | 3.2 | -27, -13, 10 | L | Putamen |
| 257 | 4.2 | -30, -22, -22 | L | Hippocampus |
| 248 | 3.3 | 11, 22, -4 | R | Caudate |
| 196 | 3.8 | -20, -37, -6 | L | Hippocampus |
| 175 | 3.4 | -10, 17, -2 | L | Caudate |
| 128 | 3.4 | -24, -13, -17 | L | Hippocampus |
| 109 | 2.9 | -17, -3, -23 | L | Amygdala |
| 92 | 2.7 | -25, -6, -14 | L | Amygdala |
| 35 | 2.6 | -1, -48, -32 | - | Brainstem |
| *PC2 × Education (+)* |  |  |  |  |
| 346 | 2.8 | -30, 2, 1 | L | Putamen |
| *PC2 × Handedness (-)* |  |  |  |  |
| 610 | 3.6 | -10, 4, 18 | L | Caudate |
| 516 | 3.3 | 10, 15, 14 | R | Caudate |
| 338 | 3.9 | -28, -12, -19 | L | Hippocampus |
| 466 | 3.7 | -23, -34, -10 | L | Hippocampus |
| 270 | 2.8 | 19, 18, 8 | R | Caudate |
| 266 | 2.5 | 19, -33, 6 | R | Thalamus |
| 231 | 2.4 | 10, -1, 8 | R | Thalamus |
| 147 | 3.7 | 28, -12, -17 | R | Hippocampus |
| 65 | 4.0 | 21, -28, -11 | R | Hippocampus |
| 39 | 2.7 | -18, -3, -16 | L | Amygdala |
| 35 | 2.4 | -18, 16, 5 | L | Caudate |
| *PC2 × Sex (-)* |  |  |  |  |
| 2507 | 3.0 | 10, -35, -28 | R | Brainstem |
| 1580 | 3.1 | 21, -10, -5 | R | Pallidum |
| 558 | 3.0 | -15, 10, 19 | L | Caudate |
| 371 | 2.7 | -23, 10, 6 | L | Putamen |
| 260 | 2.9 | -15, 1, 2 | L | Pallidum |
| 151 | 3.1 | -20, -9, -6 | L | Pallidum |
| 117 | 3.0 | -16, -7, 17 | L | Caudate |
| 107 | 3.5 | -18, 18, -3 | L | Caudate |

**Table SOM5.** Cortical results. Significant associations between variables included in the PSQI PC analysis and cortical morphometry (note that no significant associations were found between cortical morphometry and any of the variables in the PSQI total score analysis). See Fig. 5. (Abbreviations are the same as in Tab. SOM3.)

| Vol.<br>(mm<sup>3</sup>) | Cluster Peak |  |  |  |
| --- | --- | --- | --- | --- |
|  | Z-score | MNI Coord | Hem | Region |
| *PC1 × Age (+)* |  |  |  |  |
| 14371 | 3.9 | −19, −0, −21 | L | Parahipp. Gyrus |
| 1822 | 3.1 | 35, 3, 4 | R | Rolandic Operc. |
| 672 | 3.2 | −34, 20, −1 | L | Ant. Insula |
| 220 | 2.7 | 19, −52, −9 | R | Lingual Gyrus |
| 116 | 2.6 | −48, 3, −12 | L | Sup. Temp. Pole |
| 65 | 2.3 | 33, −3, −11 | R | Ant. Insula |
| 33 | 2.2 | −30, 8, 10 | L | Ant. Insula |
| 32 | 1.7 | −4, −65, 14 | L | Occipitopar. Sulc. |
| *PC1 × Age (−)* |  |  |  |  |
| 370 | 4.0 | −17, −71, 48 | L | Sup. Par. Gyrus |
| 349 | 4.3 | −33, −68, 44 | L | Intrapar. Sulc. |
| *PC2 × Education (−)* |  |  |  |  |
| 861 | 3.2 | 33, 12, 3 | R | Ant. Insula |
| 189 | 3.2 | 3, 6, −13 | R | Ant. Insula |
| *PC2 × Sex (+)* |  |  |  |  |
| 3412 | 5.1 | −2, −7, 60 | L | Supp. Motor Area |
| 852 | 4.1 | 5, 18, 41 | R | Paracing. Gyrus |

**Fig. SOM1.**
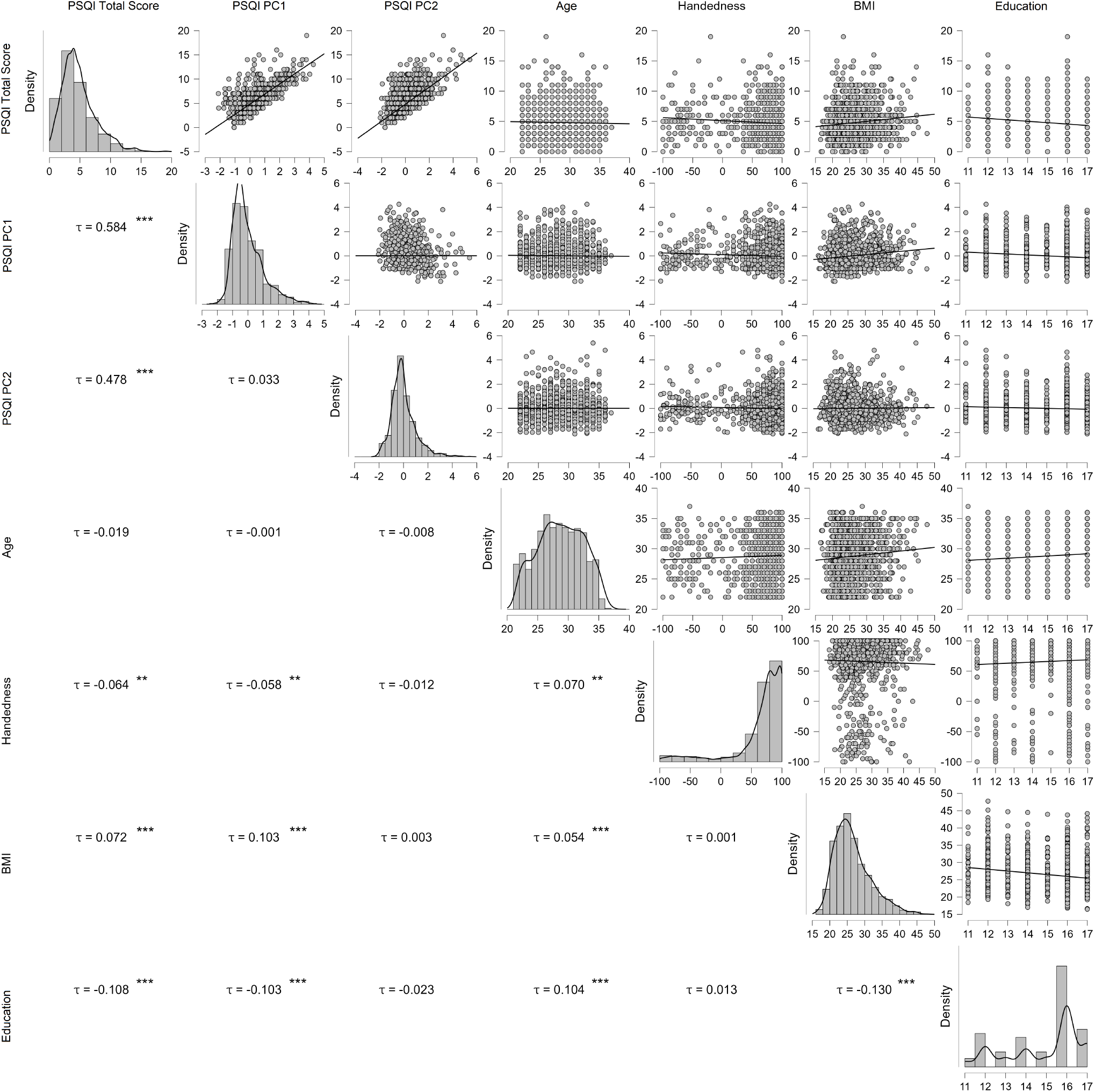
PSQI and covariates correlation structure. Pairwise correlation (Kendall’s tau () between PSQI, PSQI PC1, PSQI PC2 and covariate measures (i.e., age, handedness, BMI, years of education). (’*’ p ¡ 0.05; ‘**’ p ¡ 0.01; ‘***’ p ¡ 0.001).

**Fig. SOM2.**
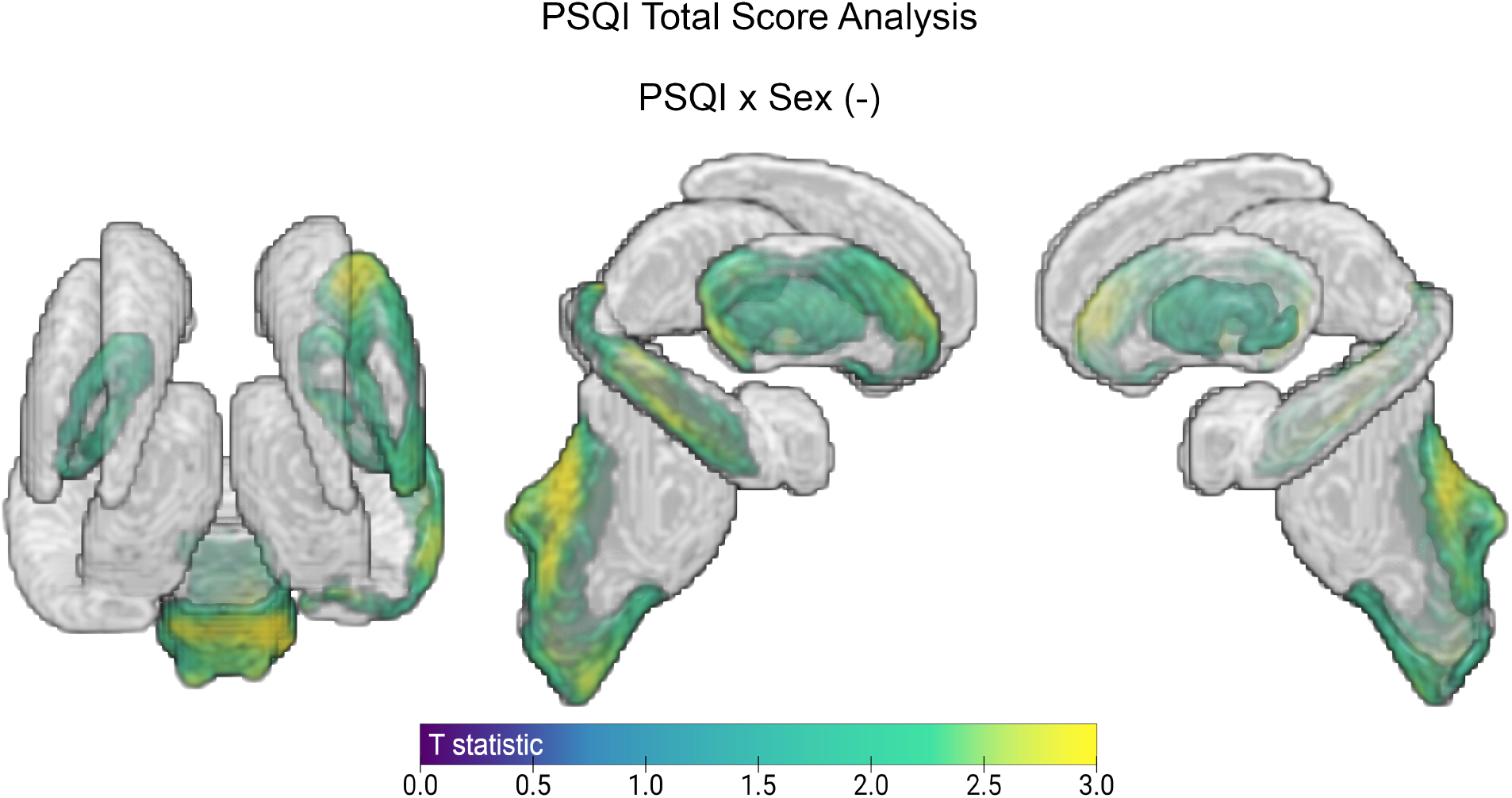
Subcortical result for the PSQI *×* Sex contrast when including only one interaction in the analysis.

